# Identification and Functional Characterization of Regulatory Variants in *DPP9* Associated with COVID-19 Severity

**DOI:** 10.1101/2025.01.07.631679

**Authors:** Gaëlle Farah, Magali Torres, Leo Henches, Hugues Aschard, Jade Ghosn, Xavier Duval, Milieu Intérieur Consortium, French COVID Cohort Study Group, Pascal Rihet, Salvatore Spicuglia, Sandrine Marquet

## Abstract

Severe acute respiratory syndrome coronavirus 2 (SARS-CoV-2) infection leads to a wide-range of clinical outcomes, which have been extensively studied through genome-wide association studies (GWAS). Starting from lead genetic variants associated with COVID-19 infection and severity, we identified a subset of non-coding candidate variants with potential regulatory functions. Using bioinformatics analysis and functional screening in three cell lines, we prioritized two *DPP9* variants within a haplotype that increases the risk of severe COVID-19. This haplotype exhibited increased regulatory activity and altered transcription factor binding, suggesting its role in influencing COVID-19 severity through modulation of DPP9 expression in immune and lung cell types. The interest of our study lies in the functional characterization of regulatory variants responsible for the higher levels of DPP9 and lung damage observed in patients with severe COVID-19. These findings advance our understanding of genetic risk factors for COVID-19 and highlight functional SNPs that may guide future therapeutic research.

## Introduction

The novel severe acute respiratory syndrome coronavirus 2 (SARS-CoV-2), the causative agent of COVID-19, was first reported in December 2019 and rapidly developed into a global pandemic, affecting almost a billion people worldwide and causing over 7 million deaths as of the 25^th^ of February 2024 according to the World Health Organization (<u>WHO Coronavirus (COVID-19) Dashboard | WHO Coronavirus (COVID-19) Dashboard With Vaccination Data)</u>. The infection causes a wide spectrum of clinical outcomes, ranging from asymptomatic carriers to individuals with severe respiratory distress and fatal outcomes. In recent years, epidemiological studies have identified several clinical risk factors for severe COVID-19, including common determinants of health and comorbidities such as hypertension, diabetes, obesity, older age, coronary artery disease, and male sex^1^. In addition to these clinical risk factors, host genetic factors have emerged as significant contributors to COVID-19 susceptibility and severity. Multiple genome-wide association studies (GWAS) have been performed to identify genetic loci associated with COVID-19 outcomes^2–4^ .Those efforts culminated with the publication of meta-analyses of several clinical phenotypes of COVID-19 across multiple populations by the COVID-19 Host Genetics Initiative (HGI). They identified a list of 13 loci associated with three different outcomes of COVID-19 (severe illness, hospitalization, and SARS-CoV-2 infection)^5^. The functional implications of these variants remain poorly understood due to a lack of comprehensive downstream analysis. Understanding the mechanistic roles of these genetic variants is crucial, as they may influence individual responses to SARS-CoV-2 infection by affecting gene expression, protein function, and immune response.

It is becoming increasingly clear that genetic variations in the non-coding genome can disrupt the finely tuned control of gene expression, leading to disease susceptibility or progression. Cis-regulatory elements can be altered in different ways that ultimately lead to dysregulation in gene expression, and therefore increase the susceptibility of the carrier individuals to a particular condition or disease. However, identifying and characterizing human variation in DNA regulatory sequences is a technically challenging task^6^. In particular, recent advances have recognized the importance of identifying causative or regulatory variants that contribute to the variability in COVID-19 clinical phenotypes^7–9^. This research seeks to deepen our understanding of the genetic underpinnings of COVID-19 and uncover novel insights into how these genetic variants may influence disease outcomes. Focusing on the 13 loci identified in the COVID-19 HGI GWAS, we investigated potential causal regulatory variants associated with COVID-19 susceptibility and severity. Combining bioinformatics approaches with experimental validation, we identified two variants that likely regulate the expression of the *DPP9* gene through epistatic interactions.

## Results

### 1 Prioritization of regulatory variants associated with COVID-19

We focused our analysis on the 13 lead variants showing a significant association with SARS-CoV-2 infection, severe manifestations of COVID-19, and hospitalization (Supplementary Table 1), as identified in the GWAS meta-analyses from Host Genetics Initiative, which included up to 49,562 European patients with COVID-19 across 19 countries^5^. First, we identified 565 SNPs in linkage disequilibrium (LD) with these lead SNPs in the European population with an r^2^ threshold of 0.6 using HaploReg V4.1^10^ and Ensembl 1000 Genome Phase 3 CEU (Fig. 1). We prioritized those variants based on their potential regulatory effect using the IW-scoring tool^11^, with detailed rankings provided in Supplementary Table 2. From this prioritized list, we selected the top 12 ranked SNPs with a K11 IW-score above 1.6 for further analysis, ensuring strong evidence of regulatory function and causality. We evaluated whether these 12 SNPs were annotated as expression quantitative trait loci (eQTL) using the GTEx browser^12^ and assessed their colocalization with ATAC-seq peaks, H3K27ac ChIP-seq peaks, and transcription factor binding sites (TFBS) using ReMap 2022^13^ (Fig. 1). Additionally, we used ENCODE^14^ data to determine whether these candidate SNPs were located in regions with potential regulatory activity, and if so, to predict whether it was preferentially a promoter or an enhancer-like region. Based on this thorough curation, 10 out of 12 SNPs were selected for downstream experimental validation. These 10 candidate SNPs were all located within H3K27ac or ATAC-seq peaks from immune cell lines^15^ (Fig. 1, supplementary Table 3), whereas the two excluded SNPs (rs74956615 and rs7306169) were neither located in regions corresponding to H3K27ac ChIP-seq or ATAC-seq peak nor annotated as a regulatory region according to ENCODE data.

**Figure 1:**
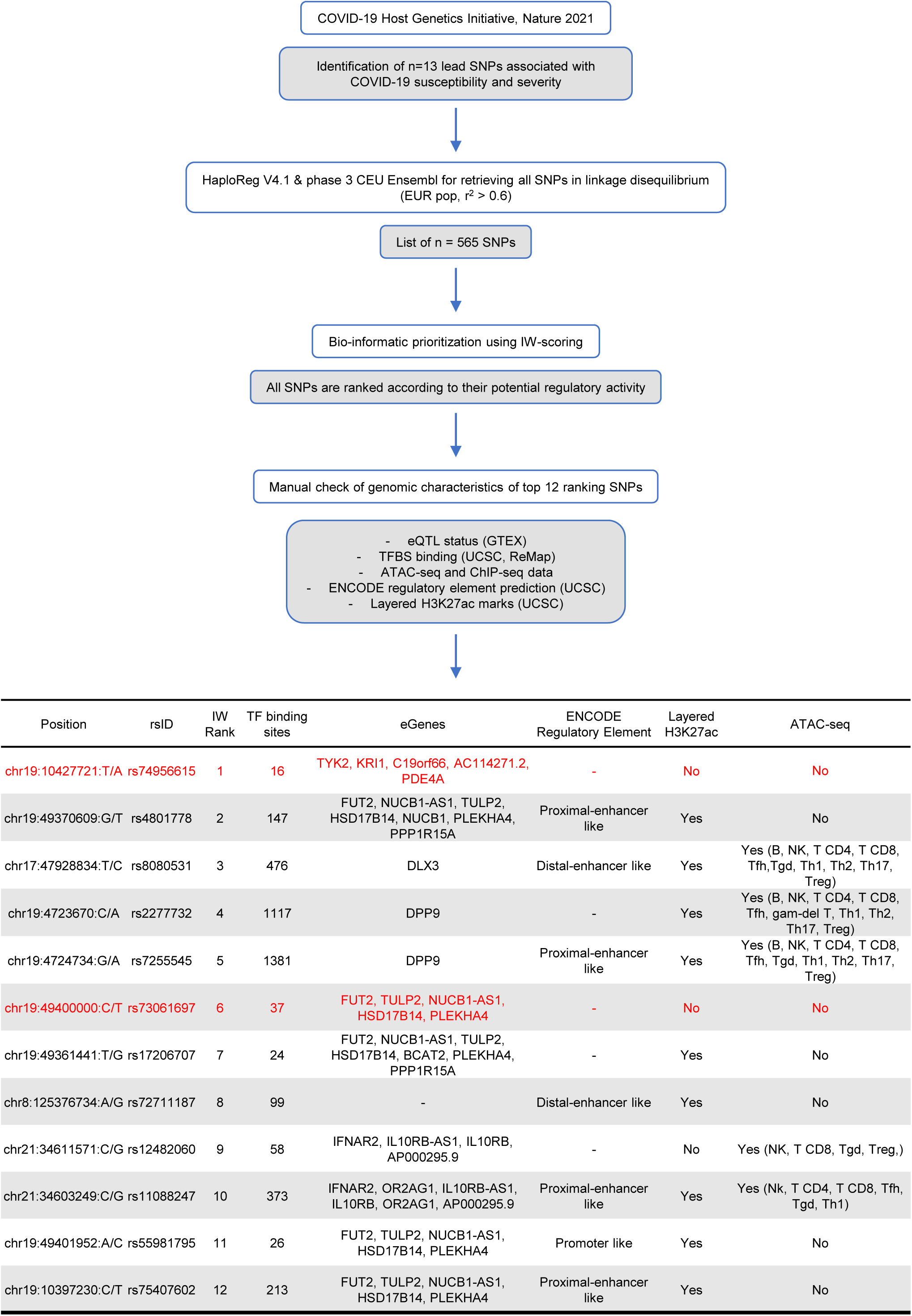
Overview of bioinformatics pipeline for prioritizing candidate causal variants at COVID-19 associated loci. From the COVID-19 HGI meta-analysis identifying 13 lead SNPs associated with COVID-19 susceptibility and severity, SNPs in linkage disequilibrium (LD) with an r^2^ threshold of 0.6 in a European population were retrieved, resulting in a list of 565 SNPs. These SNPs were prioritized using IW-scoring tool. The top 12 ranking SNPs were further examined for key genomic characteristics, including TF binding sites, target eGenes, overlap with ENCODE regulatory element predictions, H3K27ac histone modification marks, and ATAC-seq peaks. A table is provided summarizing these characteristics for each SNP. SNPs selected for downstream validation are indicated in black, while SNPs excluded from validation are highlighted in red.

### 2 Assessment of regulatory activity of candidate SNP-containing regions

Taking into consideration the genomic characteristics (distance from the TSS, annotation from ENCODE) of our 10 selected loci containing the candidate SNPs, we tested one region as a promoter (rs2277732_*DPP9*), 7 regions as potential enhancers (rs4801778_*PLEKHA4*, rs8080531_*DLX3*, rs17206707_*PLEKHA4*, rs72711187_*TMEM65*, rs12482060_*IFNAR2*, rs11088247_*IFNAR2*, rs75407602_*TULP2*) and 2 regions for both promoter and enhancer activity (rs7255545_*DPP9*, rs55981795_*TULP2*) (Fig. 2a and 2b). For each candidate variant, we selected a genomic region to test between 453 and 873 bp (Supplementary Table 3), centered around the TF density peak identified by ReMap 2022. Luciferase reporter assays were performed in two immune cell lines (K562, Jurkat) and one lung epithelial cell line (A549) with or without interferon-alpha (IFNα) stimulation to mimic a viral infection^16^ (Fig. 2a and 2b). We focused our functional analysis on these cell lines because immune cells play a major role in the host response to SARS-CoV-2, and pulmonary symptoms are the main cause of hospital admission^17^. The cells were transfected either with a construct containing the candidate regions upstream of the luciferase reporter gene to evaluate promoter function, or with a construct containing the SV40 promoter upstream of the luciferase and the candidate region downstream of the luciferase for enhancer activity. As a positive control of IFNα induction, we used an *OAS3* inducible regulatory region with both promoter and enhancer activities^18^ (Fig. 2). As shown in Figure 2a, the region rs2277732_*DPP9* containing the reference allele C of rs2277732 consistently exhibits strong promoter activity across all three cell lines compared to the basic plasmid. IFNα stimulation appears to have no effect on the tested regions in Jurkat and K562 cell lines but reduces the regulatory activity of most tested regions in the A549 cell line. No enhancer activity was observed for any of the tested regions in any of the cell lines when compared to the baseline SV40 promoter construct (Fig. 2b).

**Figure 2:**
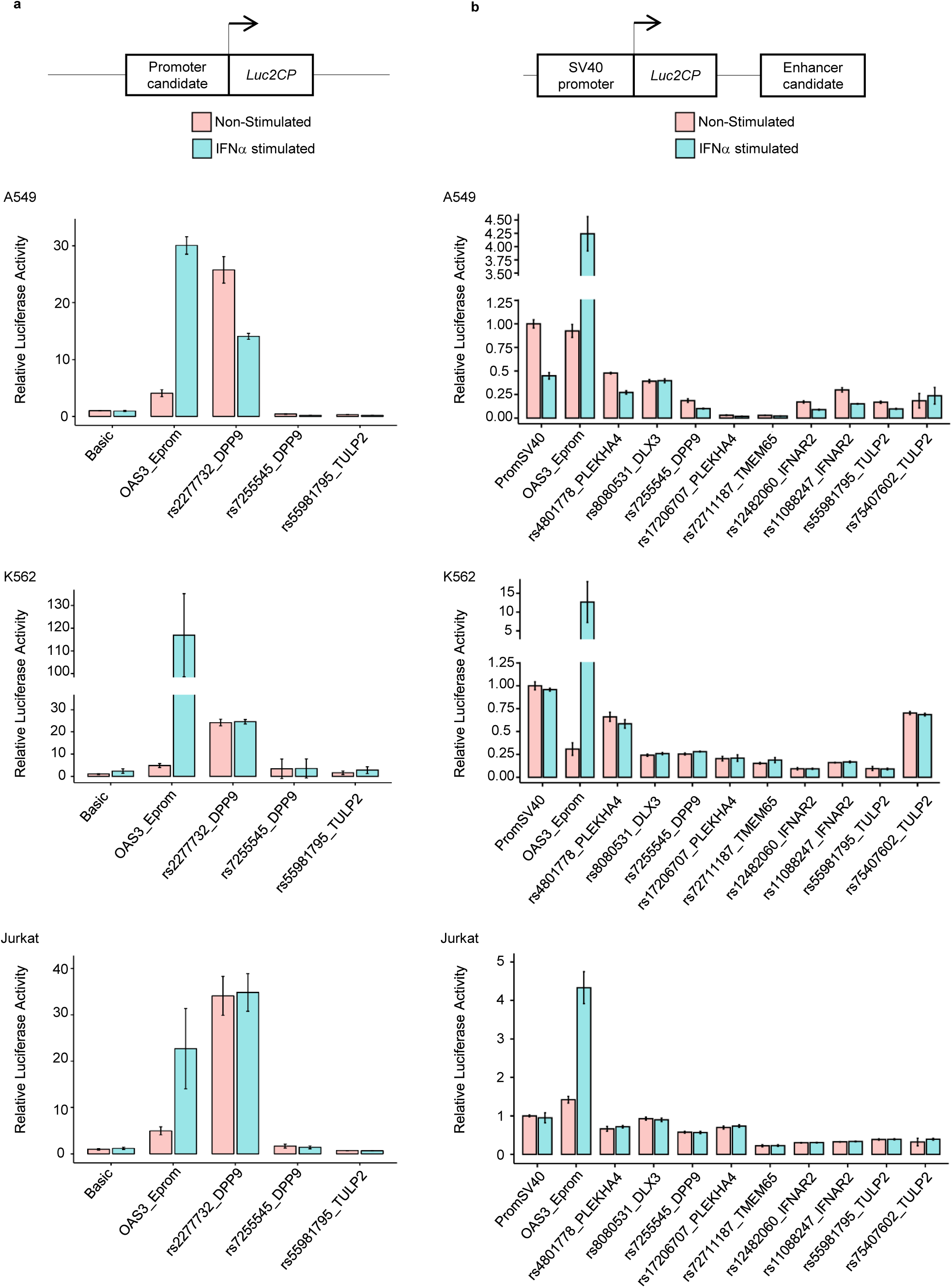
Functional screening of candidate regulatory variants across different cell lines. Luciferase reporter assay results in A549, K562 and Jurkat cell lines for regions containing the 10 selected candidate SNPs. **a** Relative luciferase activity of three regions containing rs2277732, rs7255545, and rs55981795 each, cloned as promoters upstream of *Luc2CP* gene compared to the basic plasmid. **b** Relative luciferase activity of the nine regions containing rs4801778, rs8080531, rs7255545, rs17206707, rs72711187, rs12482060, rs11088247, rs55981795 and rs75407602 each, cloned as enhancers downstream of the *Luc2CP* gene and SV40 promoter compared to the promSV40 construct. Firefly luciferase activity was measured 24h post- transfection, both without stimulation (pink bar) or following 6 hours of interferon-alpha (IFNα) stimulation (blue bar). The inducible *OAS3* Eprom construct served as positive control for IFNα stimulation. Data are represented as fold change compared to basic plasmid or promSV40. The experiments were done in triplicate.

### 3 Genetic and genomic characterization of the *DPP9* regulatory locus

Based on the luciferase assay results, we focused on *DPP9*, the gene mapped near rs2277732, for further downstream analysis. As shown in Figure 3a, the lead SNP rs2109069, previously associated with all three COVID-19 outcomes (severe illness, hospitalization, and SARS-CoV-2 infection)^5^, does not colocalize with any H3K27ac mark or ChIP-seq peaks according to the ReMap catalogue, suggesting it has no functional role. This hypothesis is further supported by the low IW-score of rs2109069, which was ranked in position 360. This SNP was found to be in LD with 6 other SNPs with r^2^>0.6 in the European population (Fig. 3a). Of these, only rs2277732 and rs7255545 had high IW scores, ranked 4^th^ and 5^th^, respectively (Supplementary Table 2). Analysis of H3K27ac profiles from the ENCODE database revealed that these SNPs are located in gene regulatory elements, with particularly strong peaks in Human lung fibroblasts (NHLF) and erythro-myeloid cells (K562) (Fig. 3a). In addition, they colocalize with ATAC-seq peaks and numerous ChIP-seq peaks, some of which correspond specifically to the binding of transcription factors in A549, K562, and Jurkat cells, the three cell lines used for the luciferase screening. According to the ENCODE data, the SNP rs7255545 is located in a cis-regulatory element with enhancer function, which appears to interact with the region containing rs2277732 (Fig. 3a). This suggests that both genetic variants may be involved in regulating the *DPP9* gene, consistent with the fact that they are both eQTLs for this gene.

**Figure 3:**
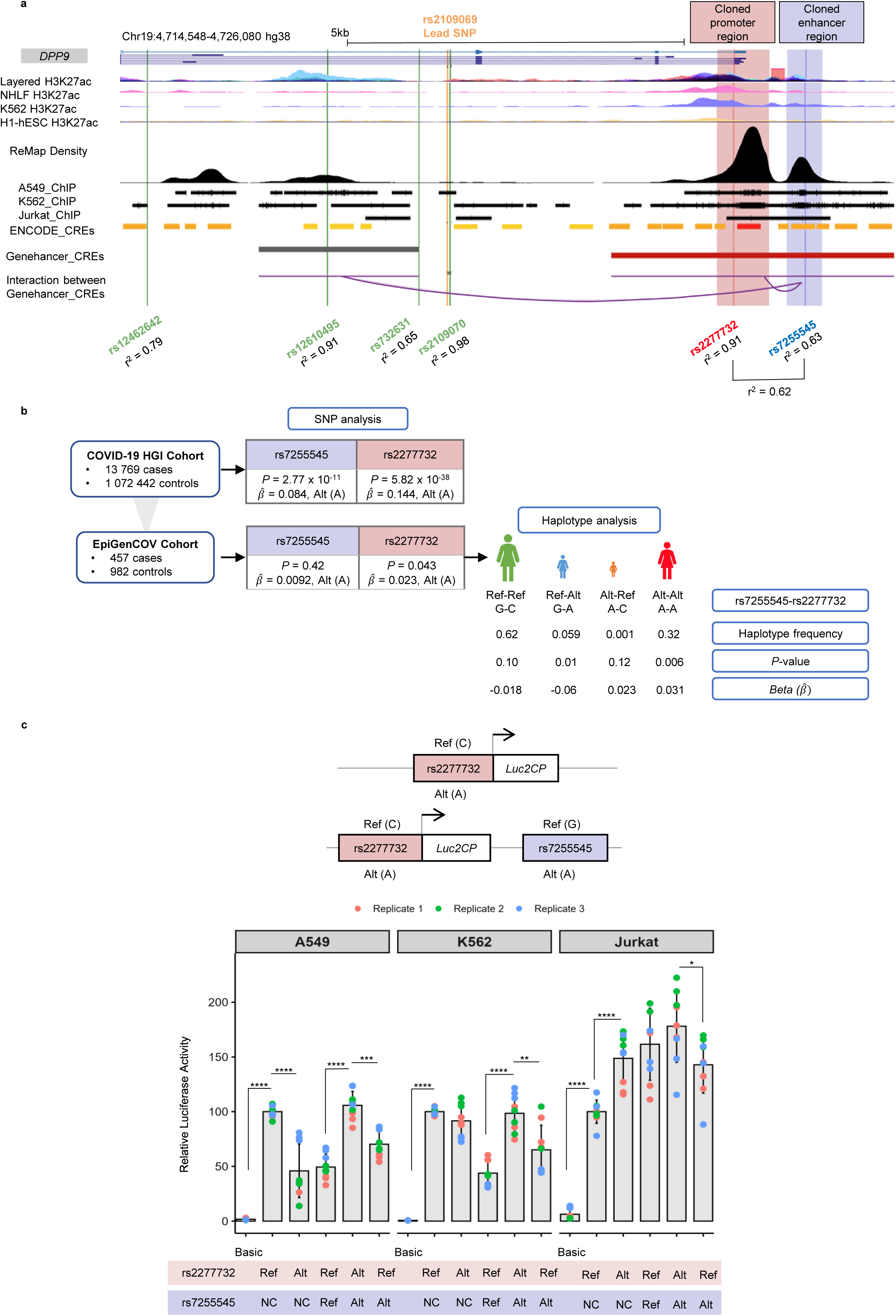
Severe COVID-19 risk haplotype in *DPP9* drives elevated gene expression. **a** Genomic characterization of the *DPP9* regulatory locus. A screenshot from the UCSC Genome Browser illustrates the genomic region surrounding the *DPP9* gene. The lead SNP rs2109069 is indicated in orange and the SNPs in linkage disequilibrium (LD) with it in green, red and blue. Our two candidate regions containing SNPs rs7255545 and rs2277732 are highlighted in blue and red respectively. The r² values for LD between rs2109069 and each SNP are indicated. The LD value between the two candidate SNPs, is also shown. Regulatory features of the genomic region are displayed, including H3K27ac histone marks across various cell types (NHLF, K562 and H1-hESC); ChIP-seq for transcription factor binding represented as ReMap density peaks, and ChIP-seq data for transcription factors in specific cell types of interest (A549, K562 and Jurkat). The screenshot also shows ENCODE CREs prediction, Genehancer CREs prediction and the interaction between Genehancer elements. **b** Genetic association of rs7255545 and rs2277732 with critical illness status in COVID-19 patients using imputation data generated from the COVID-19 HGI cohort and the EpiGenCOV cohorts. The *P*-values are indicated for both SNPs as well as the 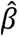 for the risk associated alternative (A) alleles. Haplotype analysis of these two variants is presented for the EpiGenCOV cohort, revealing four possible haplotypes (Ref-Ref, Ref-Alt, Alt-Ref, Alt-Alt), with their respective frequencies. The *P*-value and the 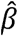 are indicated for each combination supporting association with critical illness. **c** Luciferase reporter assays of recombinant haplotypes. The constructs used for functional validation of haplotype analysis are shown. The region containing either alleles of rs2277732 (red) is cloned upstream of the *Luc2CP* gene alone (Ref-NC or Alt-NC) or with the region containing either allele of rs7255545 (blue) downstream of the *Luc2CP* gene. The different constructs were tested three times in triplicate (as indicated by the colored dots) in A549, K562, and Jurkat cell lines. Results are presented as relative luciferase activity, compared to the Ref-NC construct. Data is represented as the mean of three replicates (± SD). Two-tailed Mann-Whitney tests were performed to assess the statistical significance of the differences between conditions, with *P*-values indicated by asterisks: \**P* < 0.05, \*\**P* < 0.01, \*\*\**P* < 0.001, \*\*\*\**P* < 0.0001.

### 4 Association of *DPP9* regulatory variants with COVID-19

We first examined the association of rs2277732 and rs7255545 SNPs with COVID-19 phenotypes (susceptibility, hospitalized and critically ill) in individuals of European ancestry. The association results were pulled from the summary statistics generated by the COVID-19 Host Genetics Initiative (COVID-19 HGI) available at https://www.covid19hg.org/. Strong associations were found between rs2277732 (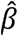 = 0.144, *P* = 5.82 x 10^-38^) and rs7255545 (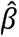 = 0.084, *P* = 2.77 x 10^-11^) and the critical illness status, indicating that the alternative A allele of each SNP is the risk allele (Fig. 3b). We also tested the association of these two SNPs in samples from EpiGenCOV, a French consortium including hospitalized COVID-19 cases from French COVID and individuals from the general population from the Milieu Intérieur cohorts^19^. We observed a nominal association with rs2277732 (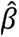 = 0.023, *P* = 0.043) but no significant association with rs7255545 (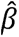 = 0.0092, *P* = 0.42) in this sub-sample of limited size, when including only one variant in the model (Fig. 3b). Interestingly, however, the joint model including the two variants produced a substantially stronger signal (*P* = 0.0047), and opposite regression coefficients (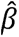*rS2277732* = 0.086 and 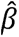*rS7255545* = -0.070), suggesting that taken together, the two variants capture a more complex structure than the sum of their individual effects.

Given the genomic proximity of SNPs rs7255545 and rs2277732, a haplotype analysis was performed to assess the combined effect of the two *DPP9* SNPs on COVID-19 critical illness. Four haplotypes were detected in the studied population, with varying numbers of individuals: ((1) *rs*7255545 **G** - rs2277732 **C**: n=1793; (2) rs7255545 **G** - rs2277732 **A:** n=148; (3) rs7255545 **A** - rs2277732 **C**: n=4; (4) rs7255545 **A** - rs2277732 **A**: n=933) (Fig. 3b). The frequency of the A-A haplotype, consisting of the alternative alleles of the two SNPs, was higher in the French COVID cohort than in the Milieu Intérieur control group. The A-A haplotype was significantly associated with a higher risk of severe COVID-19 compared to the general population (*P* = 0.006, 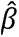 =0.031) (Fig. 3b). We also explored a homozygote model and obtained similar results (*P* = 0.005, 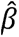 = 0.068). As for the joint analysis, the haplotype analysis strengthened the association with severe COVID-19, as compared to individual SNP analyses. This is supported by promoter-enhancer interaction data from the GeneHancer database^20^, which demonstrates interaction between the two regulatory elements containing each of these SNPs (Fig. 3a). This haplotype analysis could not be carried out in the larger cohort of COVID-19 HGI subjects, as individual data for these two SNPs, required for phasing and haplotype reconstruction, were not available.

### 5 Higher regulatory activity linked to the *DPP9* risk haplotype

To evaluate the regulatory effect of rs2277732 alleles alone or in combination with rs7255545 alleles, we performed allelic-specific luciferase reporter gene assays. First, the 769 pb region containing the rs2277732 SNP was cloned upstream of the luciferase reporter gene with either the reference C (reference) or the A (alternative) alleles to assess its allele-specific promoter function. Subsequently, we added the 524 pb region containing rs7255545 downstream of the luciferase reporter gene, as ENCODE data suggests it may act as a proximal enhancer-like region (Fig. 1). The G (reference) and A (alternative) alleles of rs7255545 were cloned with the C (reference) and A (alternative) alleles of rs2277732, respectively, to match the two most frequent haplotypes in our study population, with frequencies of 0.62 and 0.32 (Fig. 3c). We also tested the construct with the A (alternative) allele of rs7255545 with the C (reference) allele of rs2277732, which has a frequency of 0.059 in the cohort, but we did not assess the construct with the G (reference) allele of rs7255545 and the A (reference) allele of rs2277732 due to its very low frequency (0.001).

Luciferase assays were performed in A549, K562, and Jurkat cell lines. Consistent with our previous screening, the region containing rs2277732 showed strong promoter activity across all three cell lines compared to the basic plasmid (*P* < 10^-4^ by Mann-Whitney) (Fig. 3c). Specifically, the C reference allele exhibited significantly stronger activity than the A allele in A549 cells (*P* = 4.11×10□□), while the A allele was associated with higher luciferase activity in Jurkat cells (*P* = 8.22×10□□, Fig. 3c). However, luciferase expression changed when both SNP-containing regions were tested in combination. Luciferase activity was notably increased with the A–A risk haplotype (alternative alleles) compared to the C–G haplotype (reference alleles) in both the A549 lung cell line and K562 cells (*P* = 4.11×10□□). Our results indicated haplotype-specific effects, with the at-risk haplotype displaying 2.1-fold and 2.2-fold higher regulatory activity than the non-risk haplotype in A549 lung cell line and K562 hematopoietic cell line, respectively. Moreover, the protective haplotype, consisting of the C reference allele of rs2277732 and the A alternative allele of rs7255545, was associated with significantly lower expression than the at-risk haplotype across all three cell lines (A549, *P* = 1.6×10^-4^, K562, *P* = 5.67×10^-3^ and Jurkat, *P* = 1.87×10^-2^). These findings suggest that the combined effect of these SNPs, rather than a single causal variant at the *DPP9* locus, may be responsible for the observed association with severe COVID-19. The consistent association of the at-risk haplotype with the highest luciferase activity across cell lines supports the hypothesis that these SNPs are functional variants, with their combined effect likely contributing to susceptibility to severe COVID-19, as indicated by our genetic data.

### 6 The two *DPP9* regulatory variants potentially impact the binding of different transcription factors

To better understand how these two SNPs may be involved in allele-specific gene regulation, we investigated their effects on transcription factor binding sites (TFBS) in silico using RSAT tools^21^. This analysis predicted that 35 TFBS were altered by rs2277732 with *p*-value ratios ranging from 1863.64 to 10.62, including 10 activators, 17 repressors, and 8 factors with dual activity according to GeneCards (Supplementary Table 4). For rs7255545, three TFBS motifs were predicted to be modified: 1 activator, 1 repressor, and 1 with dual activity, with *p*-value ratios ranging from 11.43 to 10.88 (Fig. 4, Supplementary Table 4). As shown in Figure 4, the results for the best four *p*-value ratios indicate that the allele A for rs2277732 disrupts the binding of NFR1, HES, TCFL5 and ZBTB33. In the same way, the alternative A allele for rs7255545 inhibits the binding of MAFF and THRB, whereas it enhances the binding of MAFF. These results suggest that the molecular mechanisms underlying haplotype-specific regulation, may be partially explained by the altered transcription factor binding.

**Figure 4:**
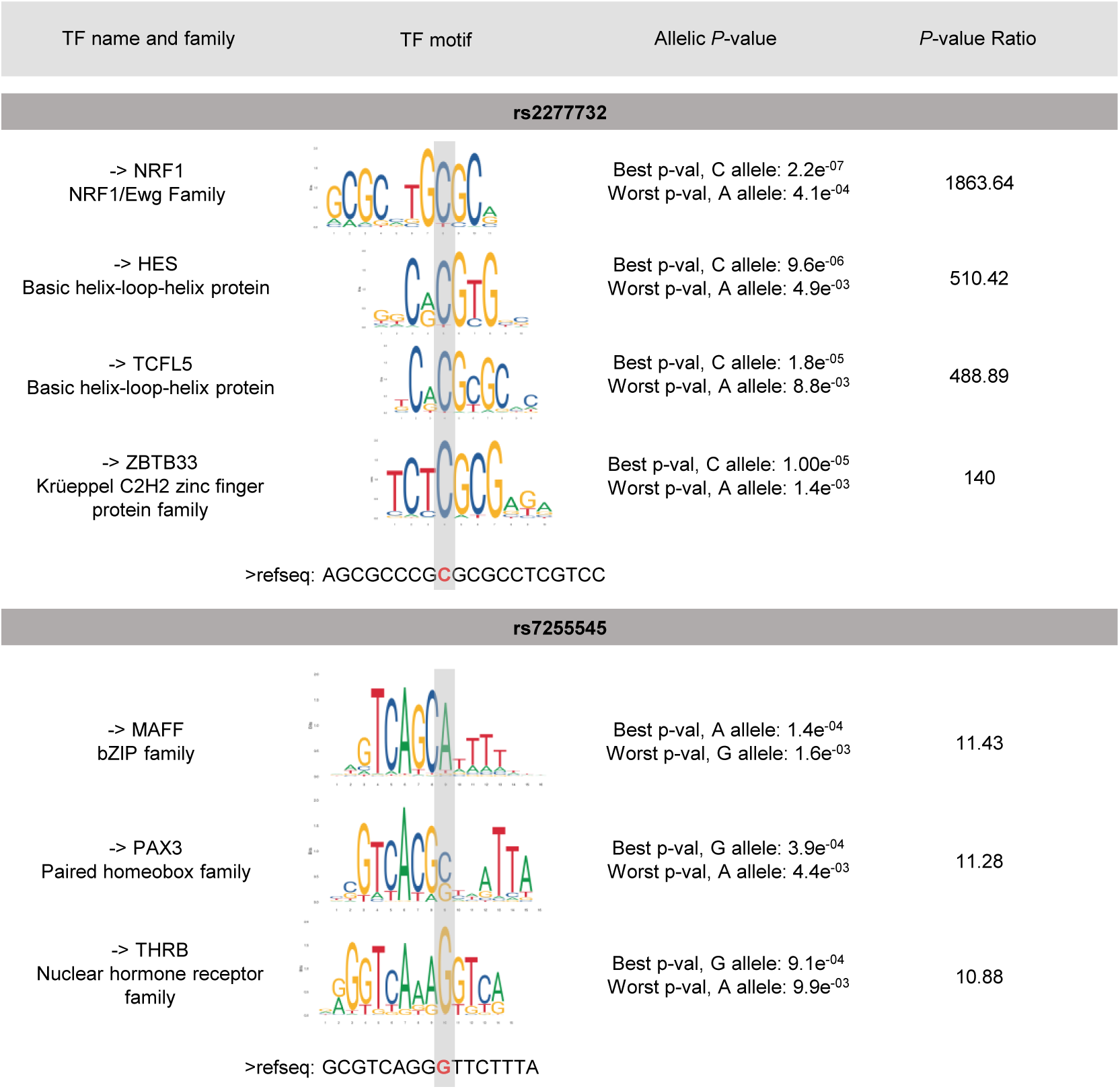
*DPP9* variants in the haplotype at-risk are predicted to disrupt the binding of several of several TFs. RSAT analysis was performed to identify the impact of the allelic configuration of rs2277732 (top) and rs7255545 (bottom) on transcription factor (TF) binding site (TFBS) motifs. The figure details the TF names and families, corresponding TF motifs and their alignment to the reference genomic sequence. Each SNP’s position within the TF motif is highlighted in grey and on the genomic sequence in red. The analysis includes the *P*-value of TF fixation for each allele for both SNPs, as well as the P-value ratio, indicating the potential disruption of TF binding due to the allelic variations.

## Discussion

Despite the availability of current vaccines and therapeutic interventions, infection with SARS-CoV-2 can still lead to severe clinical outcomes and death in some individuals. Identifying genetic factors that influence susceptibility to infection and disease severity is essential for deepening our understanding of pathophysiological mechanisms and for discovering new drug targets. GWAS and meta-analyses have provided valuable insights into the human genetic factors associated with the risk and severity of COVID-19 risk, but knowledge of the specific genes and pathways involved remains limited. A major challenge is to pinpoint the causal genes and variants responsible for these associations, particularly difficult as many of the variants are located in non-coding regions. While a few studies have investigated the potential regulatory functions of these variants through bioinformatics approaches, using eQTLs and sQTLs databases, none have provided experimental validation to clearly identify causal variants. In this study, we combined bioinformatics and experimental approaches to identify two novel causal variants that increase *DPP9* expression in lung and immune cells, thereby increasing the risk of severe COVID-19. These findings provide additional evidence for the critical role of *DPP9* in SARS-CoV2 infection and highlight its potential as a therapeutic target for preventing severe COVID-19.

We first selected the 13 lead SNPs associated with various COVID-19 phenotypes^5^ and identified 565 variants in LD with these SNPs. These variants were then prioritized based on their potential functional relevance using bioinformatics tools and epigenomic data. To further investigate the genetic regulation of the ten top-scoring variants, we performed functional analyses in three relevant cell lines. Our results showed that the haplotypic combination of the alternative alleles of two *DPP9* variants (rs2277732 and rs7255545) increases the risk of severe COVID-19 and enhances gene reporter activity in the A549 lung cell line as well as in K562 and Jurkat cell lines, compared to the non-risk haplotype, thereby suggesting increased *DPP9* expression. Moreover, we showed that as for the joint analysis, the haplotype analysis strengthened the association with severe COVID-19, compared to individual SNP analyses, supporting the combined functional effect of these two variants. Our findings are consistent with several studies reporting significantly elevated mRNA *DPP9* levels in peripheral blood of COVID-19 patients^22^ compared to healthy individuals, with levels increasing alongside disease severity^23^. The association between severe disease and *DPP9* variants, particularly rs2109069 and rs12610495, which are in LD with our candidate variants, has been replicated in multiple studies^24–26^. However, the risk alleles of both variants are paradoxically associated with reduced *DPP9* expression in the lungs, as indicated by eQTL data from GTEx. Using our pipeline, rs12610495 and rs2109069 were not prioritized by the IW scoring tool, nor did they display characteristics suggestive of potential regulatory function. Hence, our study highlights the limitations of using eQTLs alone to functionally characterize regulatory variants, as eQTLs only capture the correlation of individual SNPs with gene expression, without demonstrating a functional effect. Another key feature of GWASs is the overlap between genetic signals for COVID-19 severity and those for lung diseases, suggesting that *DPP9* may have a common role in the pathogenesis of these diseases. This is consistent with epidemiological data associating pre-existing lung conditions with more severe COVID-19 outcomes^27,28^, as well as the fact that respiratory failure remains a major cause of death among hospitalized COVID-19 patients^29,30^. Furthermore, extensive evidence indicates that pulmonary fibrosis is a hallmark of severe COVID-19^31^. This is consistent with findings that the rs12610495 variant, associated with COVID-19^32^, was also strongly associated with an increased risk of idiopathic pulmonary fibrosis (IPF) in independent studies^33–35^. In addition, higher *DPP9* expression levels have been observed in lung tissue biopsies from IPF patients compared to controls. However, neither rs12610495 nor rs2109069 were found to influence *DPP9* expression^33^, suggesting they are not causal variants. Instead, we propose that the combination of alternative alleles of the rs2277732 and rs7255545, which increase gene expression in the lung and immune cells, may contribute to the development of severe COVID-19, although the underlying mechanism remains unclear.

Recently, DPP9 has been identified as a novel repressor of the NLRP1 inflammasome activation^36^. NLRP1, highly expressed in epithelial barrier tissues, initiates the host innate immune response. Notably, human NLRP1 can detect double-stranded RNA (dsRNA) and activate an inflammasome complex in response to dsRNA detection^37^, suggesting that dsRNA replication intermediates of SARS-CoV-2^38^ could activate NLRP1, triggering an inflammatory cascade through inflammasome activation. While inflammasome activation can defend against pathogens, excessive activation may result in tissue damage and unrestrained immune cell recruitment. We propose that elevated DPP9 levels in severe COVID-19 patients may help suppress inflammation^22,23^ by inhibiting NLRP1, preventing downstream inflammasome activation^39^. Thus, DPP9 might act as an intracellular checkpoint, limiting pyroptosis and cell lysis by modulating the immune response to SARS-CoV-2. However, overexpression of DPP9 in patients with the at-risk haplotype could impair the innate immune response’s ability to restrict viral replication. This might help the virus to evade immune assault at early stages of infection and eventually leading to the release of large quantities of cytokine from dying monocytes at later stages, which would promote hyperinflammation and a cytokine storm in patients^40^.

Conversely, DPP9 also contributes to inflammation by promoting monocyte and macrophage activation. Studies indicate that DPP9 gene silencing in human monocytes^41^ or inhibition of its activity in macrophages leads to impaired activation, as indicated by reduced secretion of the proinflammatory cytokine IL-6^42^. In severe COVID-19, monocytes, and macrophages are likely major sources of the excessive proinflammatory mediators including TNF-α and IL-6 in the respiratory tract^43^. IL-6 levels have also been associated with mortality, admission to intensive care and hospitalization, representing a poor prognostic factor for COVID-19^44^. Recent reports have suggested that cytokine profiles of patients with severe COVID-19 show similarities to cytokine release syndromes such as macrophage activation syndromes^43,45,46^ and that monocytes and macrophages extensively accumulate in the lungs^47^. Positive feedback from cytokine release and subsequent immune cell recruitment can create a cycle of hyperinflammatory response, resulting in a high concentration of cytokine-secreting monocytes and neutrophils, which likely contributes to severe lung damage. Consistently, elevated *DPP9* expression has been shown to promote fibrosis in renal tubular epithelial cells^48^, and increased *DPP9* mRNA levels have been observed in inflamed lung tissues^49^. In addition, while DPP9 enzymatic activity is increased in experimental rat asthma, the mouse model of airway inflammation revealed a protective role for NLRP1 inflammasome activation by blocking DPP9^50^. Finally, it was recently shown that DPP9 also expressed in epithelial cells alters cell adhesion in human embryonic kidney cells^51^ suggesting that pulmonary fibrosis may also arise from perturbation in cell-cell adhesion or cytoskeletal integrity, which may be unable to withstand the mechanical stress of stretching within the lung. Concordant with this finding, overexpression of DPP9 suggests that it may play an important role in mediating lung damage observed in severe cases of COVID-19. Therefore, DPP9 could be used as a potential risk assessment indicator for COVID-19 patients.

In conclusion, we have identified 2 causal variants whose haplotype of alternative alleles is associated with increased risk of severe forms of COVID-19 and higher levels of DPP9. However, the function of DPP9 in SARS-CoV 2 infections appears to be complex as it may play a dual role in modulating the immune and inflammatory response. DPP9 levels may inhibit NLRP1 to suppress inflammation whereas overexpression of DPP9 may promote lung hyperinflammation by activating monocytes and macrophages as well as altering cellular integrity. All these eventually lead to a storm of cytokines and lung damage responsible for respiratory failure.

## Materials and Methods

### 1 Bio-informatic prioritization and functional annotation of genetic variants

Significant variants (n = 13) associated with different phenotypic outcomes of COVID-19 were identified as lead SNPs in the COVID-19 Host Genetics Initiative meta-analysis^5^. Using HaploReg v.4.1 <u>(broadinstitute.org)</u> and Ensembl 1000 Genome Phase 3 CEU, a list of variants of interest in LD with the 13 lead SNPs was created. HaploReg integrates regulatory genomic maps and data from ENCODE, GTEx, Roadmap Epigenomics and the 1000 Genomes Project for the purpose of mining putative causal variants, cell types, regulators and target genes for complex traits and diseases by calculating the LD between variants and assessing enrichment analysis^10^. For this analysis in a European population, with an r^2^ threshold > 0.6, we have identified 565 variants of interest. To prioritize those variants with potential functional effects, we performed a bioinformatic analysis using the IW-Scoring annotation tool (<u>SNP Annotation Tool (snp-nexus.org)</u>). This tool combines disease/phenotype and tissue/cell specificities with their related annotations into a single functional score since the tool evaluates eleven scoring methods derived from eight studies (CADD, FitCons, Eigen, FATHMM, GWAVA, DeepSEA, FunSeq2, ReMM)^11^. That way, the final score is based on diverse genomic features regarding the context of surrounding sequence gene model annotations, evolutionary constraints, epigenetic measurements, functional predictions, and assays. Furthermore, the IW-scoring tool provides two versions of its integrated score: the K10 score and the K11 score. The K11 score incorporates all 11 functional prediction tools including the FitCons tool to assess evolutionary conservation, while the K10 score excludes FitCons. In order to make sure our candidate variants are functional and are associated with causality, we employed a two-tiered thresholding approach where we only selected the variants with K11 scores >1.6 and K10 scores>1.5. This resulted in 12 candidate variants for further functional validation. We next checked if these top 12 ranking SNPs were annotated as eQTLs using GTEx browser (GTEx Portal). Moreover, using ReMap 2022 (ReMap2022 (univ-amu.fr)), we checked TF/DNA binding based on ChIP-seq experiments. To confirm whether the SNPs resided in chromatin-accessible regions, ATAC-seq data was visualized using the UCSC Genome Browser (UCSC Genome Browser). This visualization assessed whether SNPs were located within regions marked by H3K27ac, open chromatin (ATAC-seq), or predicted regulatory activity (ENCODE).

### 2 Functional screening of candidate SNPs

#### 2.1 Promoter activity

To assess promoter activity, regions ranging from 504 to 769 bp, each containing one of the following variants (rs2277732, rs7255545, or rs55981795 for the reference allele), were cloned upstream of the *Luc2CP* gene into the *NheI-BglII* site of the plasmid pGL4.12 (E6671, Promega, Madison, WI, USA). The cloning was performed by GeneCust© (GeneCust, Boynes, France).

#### 2.2 Enhancer activity

To assess enhancer activity, regions ranging from 453 to 875 bp, each containing one of the following variants (rs4801778, rs8080531, rs7255545, rs17206707, rs72711187, rs12482060, rs11088247, rs55981795 and rs75407602 for the reference allele) were cloned downstream of the *Luc2CP* gene into the *BamHI* site of the pGL4.12 plasmid containing the SV40 promoter. The cloning was performed by GeneCust© (GeneCust, Boynes, France). The pGL4.12-SV40 vector was constructed by Juliette Malfait by cloning the minimal SV40 promoter into the pGL4.12 vector (Promega, E6671) at the BglII and HindIII restriction sites.

#### 2.3 Bacterial transformation and midi prep

Lyophilized plasmids were resuspended in 10uL MilliQ de-ionized nuclease-free water, and were subsequently transformed into NEB® 10-beta electrocompetent *E.coli* DH10B strain (New England Biolabs, C3020K) by electroporation according to the manufacturer’s protocol. Transformed bacteria were cultured and harvested by centrifugation at 4000 rpm for 20 minutes, at 4 C, and plasmids were extracted using the QIAGEN Plasmid Plus Midi Kit (cat. nos. 12943 and 12945; https://www.qiagen.com/us) following the manufacturer’s instructions.

#### 2.4 Cell Culture

K562 cells (ATCC CLL-243), a chronic myelogenous leukemia cell line, and Jurkat cells (ATCC TIB-152), a T-cell acute lymphoblastic leukemia cell line, were cultured in Gibco RPMI 1640 medium (Thermo Fisher Scientific, Waltham, MA, USA), supplemented with 10% fetal bovine serum (FBS). A549 cells (ATCC CCL-185), a lung carcinoma cell line, were maintained as adherent monolayers in Dulbecco’s modified Eagle’s medium (DMEM/F12, Thermo Fisher Scientific, Waltham, MA, USA), supplemented with 10% FBS. All cell lines were grown at 37°C in a humidified incubator with 5% CO_2_ and passed every 2-3 days at a density of 3x10^5^ cells per mL. Cultures were systematically tested and confirmed to be free from mycoplasma contamination.

#### 2.5 Luciferase reporter assay

For each assay, 10^6^ K562 or Jurkat cells were co-transfected with 200ng of pRL-SV40 (a plasmid encoding Renilla luciferase) (Promega, E2231), and either 1ug of constructs of interest or 1ug of control vectors using the Neon^TM^ Transfection System (MPK5000, Thermo Fisher Scientific) according to the manufacturer’s instructions. A549 cells were co-transfected with 100ng of pRL-SV40 (Promega, E2231), and either 1ug of constructs of interest or 1ug of control vectors using Lipofectamine^TM^ 3000 Transfection Reagent kit (L3000008, Thermo Fisher Scientific). The negative control plasmid for promoter activity is an empty pGL4.12 basic and the negative control construct for enhancer activity is pGL4.12 with SV40 promoter. Sixteen hours after transfection, cells were split in two. Only half of the cells were stimulated with interferon alpha (IFNα) (SRP4594-100UG – Sigma-Aldrich). Stimulation efficiency was validated using a pGL4.12-SV40 construct with the *OAS3* Epromoter cloned downstream of the *Luc2CP* gene. This construct was generated by Antoinette Van Ouwerkerk. Firefly and Renilla luciferase activities were measured 6 hours after stimulation (24 hours post-transfection), using the Dual-Luciferase® Reporter Assay System kit (E1980, Promega), according to the manufacturer’s protocol. Measurements were performed using 20 µL of cell lysates, on the *VICTOR Nivo* machine (PerkinElmer). Firefly luciferase data were normalized to Renilla luciferase activity and expressed as fold change of relative light units over the respective negative controls.

### 3 Cohorts and genetic analysis

Individual-level haplotype data came from two cohorts assembled as part of the EpiGenCOV consortium: hospitalized COVID19 cases from the French COVID study and controls from the Milieu Interieur cohort. French COVID participants are critically ill cases of COVID-19 with COVID-19 infection confirmed by PCR and who either required respiratory support in hospital or died due to the disease. The patients were recruited across over 130 hospitals in France. Milieu Interieur controls are individual from the general population in France without known SARS-CoV-2 infection. The cohort includes over 1,000 healthy volunteers, stratified according to sex with a 1:1 ratio and age (5 decades of age spreading from 20 to 69 years old) recruited in France. For the present analysis we used a subset of 457 French COVID cases and 982 controls from Milieu Interieur, that were all European ancestry, and with both genome-wide genetic and covariates data (age, sex and body mass index).

Samples were genotyped with two different DNA arrays, and we imputed each cohort separately. Imputation was done with the RICOPILI^52^ pipeline and using the 1000 Genome reference panel. Imputation scores were 0.99 and 0.97 for rs2277732, and 0.88 and 0.92 for rs7255545 in French COVID and Milieu Interieur respectively. Relatedness between samples and population stratification was derived using chip-wide SNP data passing stringent quality control. We extracted 2 haplotypes from the imputed data for each participant. We defined 4 types of haplotypes: G-C, G-A, A-C, and A-A, depending on the observed alleles for rs2277732 (first position) and rs7255545 (second position). The sample size for each haplotype can be found in (Fig.3b). All association models included the first 20 principal components of the genotype matrix, sex, sex squared, age, and age squared as covariates. Association test was performed using a linear regression between the genetic variants of interest (either a single SNP or haplotype) and COVID-19, using the following statistical model: *Y∼*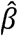*_G_G+ ∑i=1…n*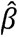*_i_C_i_*, where ci represents the covariate i. The joint test of rs2277732 and rs7255545 was performed using a multivariate Wald test of the regression coefficients 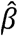*G1* and 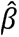*G2* from the model: *Y∼*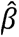*_G1_G1 +* 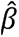*_G2_G2 + ∑i=1…n* 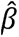*_i_C_i_*, where G1 and G2 represent rs2277732 and rs7255545, respectively.

The genotype of these two SNPs was successfully imputed in both cohorts, with imputation quality scores of 0.99 and 0.97 for rs2277732 in the case and control cohorts, respectively, and 0.88 and 0.92 for rs7255545 in the case and control cohorts, respectively.

### 5 Site-directed mutagenesis

The Q5® Site-Directed Mutagenesis Kit (New England Biolabs, E0554S) was used to substitute the reference allele (C) of the cloned SNP rs2277732 with the alternative allele (A). Mutagenesis was performed by PCR using primers designed with NEBaseChanger (https://nebasechanger.neb.com/). The primer sequences were as follows: rs2277732 (Fwd: 5’-CAGCGCCCGCTCGCCTCGTCC / Rev: 3’-CTCACCGGGACGTGGGGC). PCR products were purified using the QIAquick® PCR Purification Kit (QIAGEN, 28106) according to the manufacturer’s protocol. Mutations were validated by Sanger sequencing using primers specific to the region cloned upstream of the *Luc2CP* gene (Fwd: 5’- CATTCATTTTATGTTTCAGGTTCAGG; Rev: 3’-CTGACTGGGTTGAAGGCTCTC). Sequencing was performed by the Eurofins platform (Eurofins Genomics - Genomic services by experts) and chromatograms were analyzed with SnapGene Viewer (https://www.snapgene.com/).

### 6 Gibson assembly

Plasmids with the reference and alternative alleles of rs2277732 were used to generate haplotype combinations (Alt/Alt, Ref/Ref and Ref/Alt) using the Gibson assembly method (Gibson Assembly® Master Mix – Assembly (E2611) NEB) following the manufacturer’s instructions. The fragment containing the reference allele (G) of rs7255545 was first retrieved by BamHI enzymatic digestion of the construct pGL4.12-SV40., purified on gel using the PureLink® Quick Gel Extraction Kit (Invitrogen, K2100-12) and cloned in the BamHI site downstream of the *Luc2CP* gene in the construct containing either allele of rs2277732. Cloning was validated by Sanger sequencing using primers that target the regions downstream of Luc2CP gene (Fwd: 5’- AGTGCAGGTGCCAGAACATT; Rev: 3’-GCCTTATGCAGTTGCTCTCC). Using the Ref-Ref and Alt-Ref constructs, we generated the Ref-Alt and Alt-Alt constructs using the Q5 site directed mutagenesis with the following primers targeting rs7255545 (Fwd: 5’- GGAGAATTTAAAGAAGCCTGA; Rev: 3’-TCAGGCTTCTTTAAATTCTCC). Mutations were validated by Sanger sequencing with the same primers used to validate Gibson cloning. The resulting constructs were amplified and transfected into K562, Jurkat, and A549 cells as described previously. Luciferase assays were performed as described above three times in triplicate.

### 7 Transcription Factor motif analysis

The impact of the selected variants on transcription factor binding sites (TFBS) was evaluated using the RSAT tool^21^ and the JASPAR database^53^.

### 8 Statistical tests

Statistical significance between luciferase constructs was assessed using a two-tailed non-parametric Wilcoxon-Mann-Whitney test performed using Rstudio (version 4.4.0). This test was applied to validate the results of the allele specific luciferase assays. Firefly luciferase data was normalized to Renilla Luciferase data, and the fold change was evaluated proportionally to the negative control (pGL4 basic). Statistical significance is indicated in the figures, with asterisks denoting *P*-values as follows: *P*< 0.05 (*), *P*< 0.01 (**), *P*< 0.001 (***), and *P*< 0.0001 (***).

## Data availability

All other data are available in the article and its Supplementary files or from the corresponding author upon request.

## Acknowledgments

We thank the CNRS (Centre national de la recherche scientifique) for providing Sandrine Marquet’s salary. This work was supported by the recurrent funding from Institute National de la Santé et de la Recherche Médicale (INSERM), Aix-Marseille University and grants from the French Agency for Research (Agence Nationale de la Recherche ANR) ANR-21-CO14-0001-01 and ANR-23-CE12-0008-01 grants. We thank the members of TAGC, Juliette Malfait and Antoinette Van Ouwerkerk for providing the pGL4.12-SV40 and pGL4.12-SV40-OAS3 plasmids respectively. Gaëlle Farah was supported by funding from LIGUE contre le Cancer. This project was carried out in the framework of the Inserm GOLD Cross-Cutting program.

We thank all members of the consortia listed below:

Members of French COVID Cohort Study Group: Amal Abrous^1^, Delphine Bachelet^2^, Marie Bartoli^3^, Lila Bouadma^2^, Minerva Cervantes-Gonzalez^2^, Anissa Chair^2^, Charlotte Charpentier^2^, Sandrine Couffin-Cadiergues^1^, Nathalie De Castro^4^, Diane Descamps^2^, Hang Doan^2^, Céline Dorival^5^, Xavier Duval^2^, Hélène Esperou^1^, Aline-Marie Florence^2^, Jade Ghosn^2^, François Goehringer^6^, Maxime Goyat^2^, Ikram Houas^1^, Isabelle Hoffmann^2^, Salma Jaafoura^1^, Simon Jamard^7^, Nadhem Lafhej^2^, Cédric Laouénan^2^, Soizic Le Mestre^8^, France Mentré^2^, Christelle Paul^8^, Aurélie Papadopoulos^1^, Marion Schneider^2^, Coralie Tardivon^2^, Sarah Tubiana^2^, Aurélie Wiedemann^9^

^1^Inserm sponsor, Paris, France. ^2^Hôpital Bichat, Paris, France. ^3^ANRS, Paris, France. ^4^Hôpital Saint Louis, Paris, France. ^5^Inserm UMR 1136, Paris, France. ^6^CHU, Nancy, France. ^7^Hôpital Bretonneau, Tours, France. ^8^ANRS-MIE, Paris, France. ^9^Vaccine Research Institute (VRI), Inserm UMR 955, Créteil, France.

This work benefited from the support of the French government’s Invest in the Future programme. This programme is managed by the Agence Nationale de la Recherche, reference ANR-10-LABX-69-01.

The Milieu Intérieur Consortium (unless otherwise indicated, partners are located at Institut Pasteur, Paris) is composed of the following team leaders: Laurent Abel^1^, Andres Alcover^2^, Hugues Aschard^2^, Philippe Bousso^2^, Nollaig Bourke^2^, Petter Brodin^3^, Pierre Bruhns^4^, Nadine Cerf-Bensussan^4^, Ana Cumano^5^, Christophe D’Enfert^5^, Caroline Demangel^5^, Ludovic Deriano^5^, Marie-Agnès Dillies^5^, James Di Santo^5^, Gérard Eberl^5^, Jost Enninga^5^, Jacques Fellay^5^, Ivo Gomperts-Boneca^3^, Milena Hasan^3^, Gunilla Karlsson Hedestam^3^, Serge Hercberg^6^, Molly A Ingersoll^7^, Olivier Lantz^8^, Rose Anne Kenny^2^, Mickaël Ménager^4^, Hugo Mouquet^2^, Cliona O’Farrelly^2^, Etienne Patin^4^, Antonio Rausell^4^, Frédéric Rieux-Laucat^4^, Lars Rogge^9^, Magnus Fontes^9^, Anavaj Sakuntabhai^10^, Olivier Schwartz^10^, Benno Schwikowski^10^, Spencer Shorte^10^, Frédéric Tangy^10^, Antoine Toubert^10^, Mathilde Touvier^6^, Marie-Noëlle Ungeheuer^11^, Christophe Zimmer^11^, Matthew L. Albert^11^ (Octant), Darragh Duffy (co-coordinator of the *Milieu Intérieur* Consortium), Lluis Quintana-Murci (co-coordinator of the *Milieu Intérieur* Consortium).

^1^Hôpital Necker. ^2^Trinity College Dublin. ^3^Karolinska Institutet. ^4^INSERM UMR 1163 – Institut Imagine. ^5^EPFL, Lausanne. ^6^Université Paris 13. ^7^Institut Cochin and Institut Pasteur. ^8^Institut Curie. ^9^Institut Roche. ^10^Hôpital Saint-Louis. ^11^Octant.

Additional information can be found at: http://www.milieuinterieur.fr

## Authors’ contributions

G.F., P.R., S.S., and S.M. performed bioinformatic analysis. L.H. and H.A. performed imputation and association studies. G.F. and M.T. performed experiments. G.F. analyzed experimental data and provided figures and tables. J.G. and X.D. participated in the recruitment of the French Covid cohort. S.S. and S.M. supervised the project. G.F. and S.M. wrote the paper. All the authors edited the manuscript. All authors read and approved the final manuscript.

## Supplementary material and information

### Milieu Interieur cohort

The Milieur Intérieur Cohort was approved by the Comité de Protection des Personnes – Ouest 6 (Committee for the protection of persons) on June 13th, 2012 and by French Agence nationale de sécurité du médicament (ANSM) on June 22nd, 2012. The study is sponsored by Institut Pasteur (Pasteur ID-RCB Number: 2012-A00238-35), and was conducted as a single centre interventional study without an investigational product. The original protocol was registered under ClinicalTrials.gov (study# NCT01699893). The samples and data used in this study were formally established as the Milieu Interieur biocollection (NCT03905993), with approvals by the Comité de Protection des Personnes – Sud Méditerranée and the Commission nationale de l’informatique et des libertés (CNIL) on April 11, 2018.

**Supplementary PDF file 1, containing supplementary Table 1 and Table 3**.

**Supplementary Table 2, Excel file containing additional data too large to fit in a PDF file.**

**Supplementary Table 4, Excel file containing additional data too large to fit in a PDF file.**

## Supplementary titles and legends

Supplementary Table 1: Coordinates, SNP ID and alleles of the 13 lead SNPs identified in the COVID19 HGI meta-analysis.

Supplementary Table 2: IW scoring tool output; coordinates and scores of 565 identified SNPs according to severals tools employed by IW scoring (CADD, DeepSea, EIGEN, FATHMM, fitCons, funSeq, GWAVA and REMM), ranked by overall K11 and K10 scores with SNP IDs.

Supplementary Table 3: Genomic characteristics of the 10 selected candidates SNPs (coordinates, SNP ID and alleles, TF binding density, ENCODE CREs, eGenes, H3K27ac marks and ATAC-seq peak overlap), as well as the associated COVID-19 phenotype of their respective lead SNP according to the COVID19 HGI meta-analysis, the tested effect for the cloned region containing each SNP and the coordinated of the selected region for the luciferase assay.

Supplementary Table 4: RSAT results showing the predicted TF motifs affected by the allelic configuration of rs2277732 and rs7255545.

