## Supplementary Table 1 and Table 3 for "Identification and Functional Characterization of Regulatory Variants in *DPP9* Associated with COVID-19 Severity"

| SNP coord | SNP iD |
|---|---|
| chr3:45796521 | rs2271616 (G>A/T) |
| chr3:101705614 | rs11919389 (T>C) |
| chr3:45823240 | rs10490770 (T>C) |
| chr6:41534945 | rs1886814 (A>C) |
| chr8:124324323 | rs72711165 (T>C) |
| chr9:133274084 | rs912805253 (C>T) |
| chr12:112919388 | rs10774671 (G>A) |
| chr17:46142465 | rs1819040 (T>A) |
| chr17:49863303 | rs77534576 (C>T) |
| chr19:4719431 | rs2109069 (G>A) |
| chr19:10317045 | rs74956615 (T>A) |
| chr19:48867352 | rs4801778 (G>T) |
| chr21:33242905 | rs13050728 (T>C) |

Supplementary Table 1

| SNP coord | SNP iD | SNP characteristics | Associated phenotype of tagSNP in LD | Tested effect | Cloned region coord |
|---|---|---|---|---|---|
| chr19:48867352 | rs4801778 (G>T) | 147 TFBS, 7 eGenes, ENCODE proximal enhancer-like region, overlaps layered H3K27ac marks | reported SARS-CoV-2 infection | Enhancer | chr19:48867010-48867536 |
| chr17:49851471 | rs8080531 (T>C) | 476 TFBS, 1 eGene (DLX3), ENCODE distal-enhancer like region, overlaps layered H3K27ac marks | critically ill SARS-CoV2-positive | Enhancer | chr17:49851184-49852025 |
| chr19:4723658 | rs2277732 (C>A) | 1117 TFBS, 1 eGene (DPP9), overlaps layered H3K27ac marks | reported SARS-CoV-2 infection, hospitalized SARS-CoV-2 positive, critically ill SARS-CoV-2 positive | Promoter | chr19:4723422-4724190 |
| chr19:4724722 | rs7255545 (G>A) | 1381 TFBS, 1 eGene (DPP9), ENCODE proximal-enhancer like region, overlaps layered H3K27ac marks | reported SARS-CoV-2 infection, hospitalized SARS-CoV-2 positive, critically ill SARS-CoV-2 positive | Promoter & enhancer | chr19:4724448-4724971 |
| chr19:48858184 | rs17206707 (T>G) | 24 TFBS, 7 eGenes, overlaps layered H3K27ac marks and ATAC-seq peaks | reported SARS-CoV-2 infection | Enhancer | chr19:48857999-48858451 |
| chr8:124364493 | rs72711187 (A>G) | 99 TFBS, ENCODE distal enhancer-like region, overlaps layered H3K27ac marks and ATAC-seq peaks | hospitalized SARS-CoV-2 positive | Enhancer | chr8:124363770-124364642 |
| chr21:33239266 | rs12482060 (C>G) | 58 TFBS, 4 eGenes, overlaps ATAC-seq peaks | hospitalized SARS-CoV-2 positive, critically ill SARS-CoV-2 positive | Enhancer | chr21:33238925-33239468 |
| chr21:33230944 | rs11088247 (C>G) | 373 TFBS, 6 eGenes, ENCODE proximal enhancer-like region, overlaps layered H3K27ac marks | hospitalized SARS-CoV-2 positive, critically ill SARS-CoV-2 positive | Enhancer | chr21:33230625-33231331 |
| chr19:48898695 | rs55981795 (A>C) | 26 TFBS, 5 eGenes, ENCODE promoter-like region, overlaps layered H3K27ac marks | reported SARS-CoV-2 positive | Promoter & enhancer | chr19:48898439-48898942 |
| chr19:10286554 | rs75407602 (C>T) | 213 TFBS, 5 eGenes, ENCODE proximal enhancer-like region, overlaps layered H3K27ac marks | hospitalized SARS-CoV-2 positive, critically ill SARS-CoV-2 positive | Enhancer | chr19:10286354-10287050 |

Supplementary Table 3
